# Flexible Skull-Conformal Phased Array for Aberration-Corrected Transcranial Focused Ultrasound Therapy

**DOI:** 10.1101/2025.11.28.691082

**Authors:** Jiayi Zhang, Haiguo Qin, Cong Pu, Ting Zhang, Junfeng Sun, Chang Peng

**Author notes:** Corresponding author: Chang Peng, Address: School of Biomedical Engineering, 393 Middle Huaxia Road, Pudong, Shanghai, 201210, China.

## Abstract

Transcranial focused ultrasound (tFUS) is a promising non-invasive tool for precise deep-brain interventions, including neuromodulation, blood-brain barrier opening, and drug delivery. However, conventional rigid tFUS arrays suffer from poor skull conformity, compromising patient comfort and targeting precision, while skull-induced aberrations degrade focal accuracy and efficiency. To address these challenges, we present the first flexible, skull-conformal aberration-corrected phased array (F-SCAPA), a transformative solution integrating mechanical adaptability with advanced acoustic correction. The 0.5-MHz array—comprising 8×8 elements—features an optimized fabrication process that enhances material efficiency, ensures flexibility and stretchability, and maintains uniform electrical performance. Coupled with CT-based skull modeling and time-reversal aberration correction, F-SCAPA dynamically conforms to cranial anatomy while compensating for phase distortions. Transcranial validation in a human skull demonstrates precise focusing at 36 mm depth, achieving an axial full width at half maximum (FWHM) of 8.9 mm. This breakthrough overcomes existing limitations by combining mechanical adaptability with high-resolution transcranial focusing, establishing F-SCAPA as a robust, patient-compatible platform for next-generation non-invasive brain therapies.

## 1. Introduction

Transcranial focused ultrasound (tFUS) is a versatile non-invasive technology for delivering acoustic energy to deep-brain targets with millimeter-scale precision^1^. Its effects are determined by acoustic parameters, enabling mild thermal activation of thermo-sensitive ion channels and mechanical stimulation of mechanosensitive channels to modulate neuronal activity^2–4^. In addition, Ultrasound-driven cavitation can also transiently permeabilize cell membranes, facilitating ion transport and therapeutic delivery across the blood–brain barrier^5,6^. Compared with electrical or magnetic stimulation, tFUS can access deep-brain structures without direct implantation, reducing tissue damage and inflammatory responses with invasive procedures^7,8^. Its ability to focus acoustic energy through the skull allows spatially selective interventions, providing opportunities for neuromodulation, transient BBB opening, and localized therapeutic delivery^9^.

Despite the therapeutic potential of transcranial focused ultrasound, existing devices face significant challenges in achieving precise targeting while maintaining patient comfort. The skull’s heterogeneous structure causes acoustic attenuation and phase aberration, leading to focal distortion, positional shifts, and energy scattering^10^. Individual differences in skull thickness, density, and curvature further complicate ultrasound delivery, reducing energy transmission efficiency and reproducibility, and requiring patient-specific alignment and calibration to achieve reliable therapeutic outcomes^11^. Traditional tFUS transducers are typically rigid and bulky, preventing them from conforming to the complex curvature of the head^12^. Their fixed focal lengths also limit the ability to target multiple brain regions without repositioning or replacing the device, reducing treatment versatility. As a result, external fixation systems are often necessary to maintain proper alignment and acoustic coupling, which increases setup complexity and compromises patient comfort and mobility^13,14^. These limitations compromise focusing accuracy and restrict treatment flexibility, hindering the development of long-term, patient-friendly transcranial ultrasound systems.

In recent years, extensive efforts have focused on developing advanced tFUS transducers. Legon et al. employed a 0.5 MHz single-element transducer to modulate the human primary somatosensory cortex^15^, demonstrating millimeter-scale modulation of cortical excitability and providing early evidence for non-invasive ultrasound-based human brain treatment, although spatial targeting flexibility remained limited. To increase flexibility and multi-region targeting, array-based transducers have been explored in previous studies. Kim et al. designed a miniaturized, head-mountable 32-element ring array (183 kHz) for freely moving mice, demonstrating motor responses and enabling in vivo experiments beyond traditional stereotaxic fixation^16^. Li et al. proposed a 128-channel linear array with a 5 MHz center frequency for image-guided neuromodulation, showing that stimulation of different focal points evoked distinct forelimb movements and electromyography (EMG) responses^17^. Zhang et al. developed a 1 MHz, 256-element 2-D array capable of achieving multi-target blood–brain barrier (BBB) opening in rats, though without evaluation through human skulls^18^.

Despite these advances, translating array-based tFUS systems to humans remains challenging due to the skull’s complex acoustic properties, which reduce focusing accuracy and necessitate precise transducer positioning and design optimization. Adams et al. implemented a skull-conformal 4096-element phased array integrated into a 3D-printed helmet scaffold optimized for ex vivo human skull caps, which validated for transcranial focusing and safe BBB opening in rodents^19^. While this approach achieved precise targeting, it was time-consuming, costly, and required individualized fabrication. Manuel et al. developed a 1 MHz, 128-element Fermat-spiral array capable of transcranial focusing without subject-specific customization and offering wide-range beamforming capability^20^. Martin et al. designed a 256-element helmet-shaped array (555 kHz) with stereotactic positioning, MRI/CT-based planning, and real-time fMRI compatibility, achieving robust theta-burst neuromodulatory effects in humans^21^. However, current tFUS transducers remain rigid and bulky, require complex acoustic coupling, and pose challenges for patient comfort and clinical usability, underscoring the need for flexible, user-friendly transcranial ultrasound systems.

In this study, we present a flexible skull-conformal aberration-corrected phased array (F-SCAPA) that integrates three significant advancements: (1) a streamlined fabrication process offering reduced complexity while maintaining high performance, (2) enhanced mechanical flexibility enabling reliable conformity to diverse cranial anatomies with high element-to-element uniformity across the 64-element array, and (3) an integrated 3D array-camera localization system coupled with CT-derived skull modeling for personalized aberration correction. The F-SCAPA system eliminates the need for individualized array redesign while establishing precise focusing capabilities validated through simulations and experiments. Compared to existing systems, our platform significantly reduces setup time and fabrication complexity, while demonstrating versatile clinical applicability for neuromodulation, BBB opening, and targeted therapy delivery. This combination of simplified implementation and advanced performance positions the F-SCAPA as a practical solution for both neuroscience research and clinical neurotherapy applications.

## 2. Results

### 2.1 Device Design and Structural Optimization

Fig. 1a schematically illustrates the design and working principle of the F-SCAPA. The device integrates a flexible 8 × 8 PZT-4 piezoelectric array for focused ultrasound generation. To achieve high acoustic pressure with low driving voltage, PZT-4 was selected as the piezoelectric material due to its high electromechanical efficiency^22,23^ (Supplementary Table. 1). For transcranial applications, a central frequency of 0.5 MHz was selected, as it offers a good balance between penetration depth and focus precision while maintaining relatively low skull-induced attenuation^24–26^. Ultrasound gel served as the coupling medium between the F-SCAPA and the skull to ensure efficient acoustic transmission. During operation, the array was secured to the skull with medical tape to prevent movement. The spatial position of each element was captured by a 3D array-camera and co-registered with the CT-derived skull model. The target region was defined within the transcranial simulation model, which calculated the optimal time delays for all 64 channels to achieve precise acoustic focusing. After parameter verification, the Vantage HIFU system was used to drive the array, generating and steering the focused ultrasound beam toward the target. The focused ultrasound could produce thermal, mechanical, or cavitation effects, enabling various transcranial therapeutic applications.

**Fig. 1.**
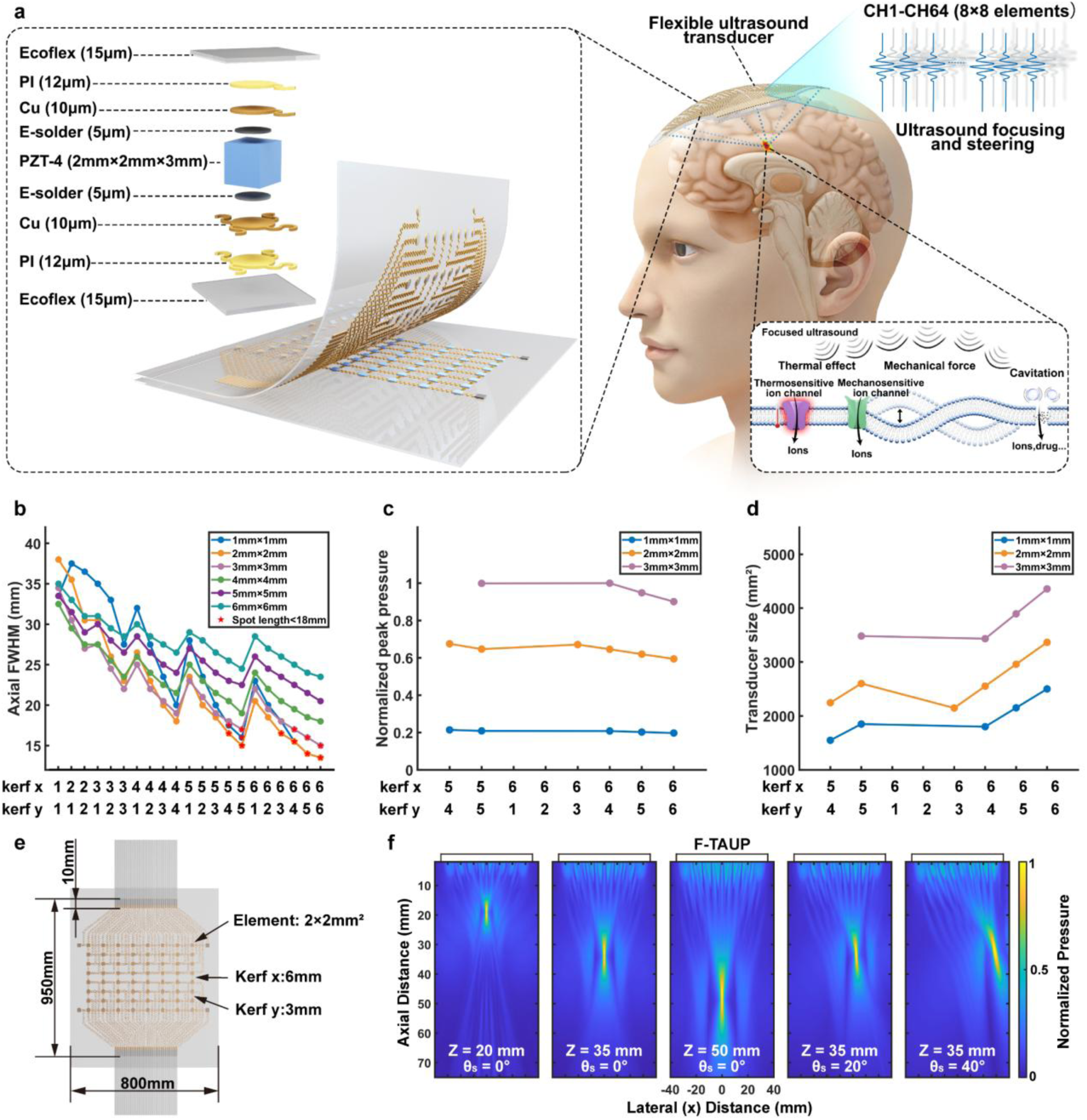
Design and working principle of the F-SCAPA. **(a)** Schematic of the operating principle. **(b)** Simulated axial FWHM for 8 × 8 arrays with different element sizes and kerfs. **(c-d)** Designs achieving axial FWHM below 18 mm: **(c)** normalized peak pressure and **(d)** transducer dimensions. **(e)** Final structural design of the array. **(f)** Simulated acoustic beams focused at depths of 20, 35, and 50 mm and steered at 20° and 40° (35 mm depth).

For transcranial applications, the F-SCAPA was designed as a two-dimensional phased array, enabling dynamic beamforming for three-dimensional focusing^27^. This configuration allows simultaneous generation of multiple foci or real-time adjustment of focal depth and steering angle, with each element individually phase-corrected to precisely control the acoustic field. Field II simulations were conducted to evaluate the acoustic performance of arrays with different structural parameters. Since therapeutic targets are typically located 20-50 mm beneath the skull^28^, a representative focal depth of 35 mm was selected for simulations. The axial Full Width at Half Maximum (FWHM) was used as the primary criterion for optimization. Arrays with element sizes ranging from 1 × 1 mm² to 6 × 6 mm² and kerf widths from 1 to 6 mm in both directions were systematically evaluated to determine the optimal design parameters.

The resulting axial FWHM values for all designs are shown in Fig. 1b, and designs achieving axial FWHM below 18 mm are summarized in Fig. 1c–d. Among these, an array with 2 × 2 mm² elements and kerfs of 6 mm (x-direction) and 3 mm (y-direction) offered the best compromise between peak acoustic pressure and a compact aperture suitable for transcranial applications. The final F-SCAPA design is illustrated in Fig. 1e, and simulations of its acoustic fields (Fig. 1f) demonstrate effective focusing and beam-steering capabilities, achieving focal points at depths of 20, 35, and 50 mm, with steering angles of 20° and 40° at 35 mm depth.

Following the fabrication process illustrated in Fig. 2, the F-SCAPA was produced by directly spin-coating a silicone elastomer (Ecoflex 00-30) onto polyvinyl chloride (PVC) substrates, followed by electrode patterning on the Ecoflex surface. Preliminary experiments confirmed that this approach enable effectively removed excess material while retaining the desired electrode patterns (Supplementary Fig. 1). This optimized, transfer-print-free process enabled fabrication of the flexible ultrasound transducer with a streamlined workflow and shorter preparation time. The simplified procedure and improved material utilization decrease labor and fabrication time, offering a feasible pathway toward scalable production and potential clinical translation.

**Fig. 2.**
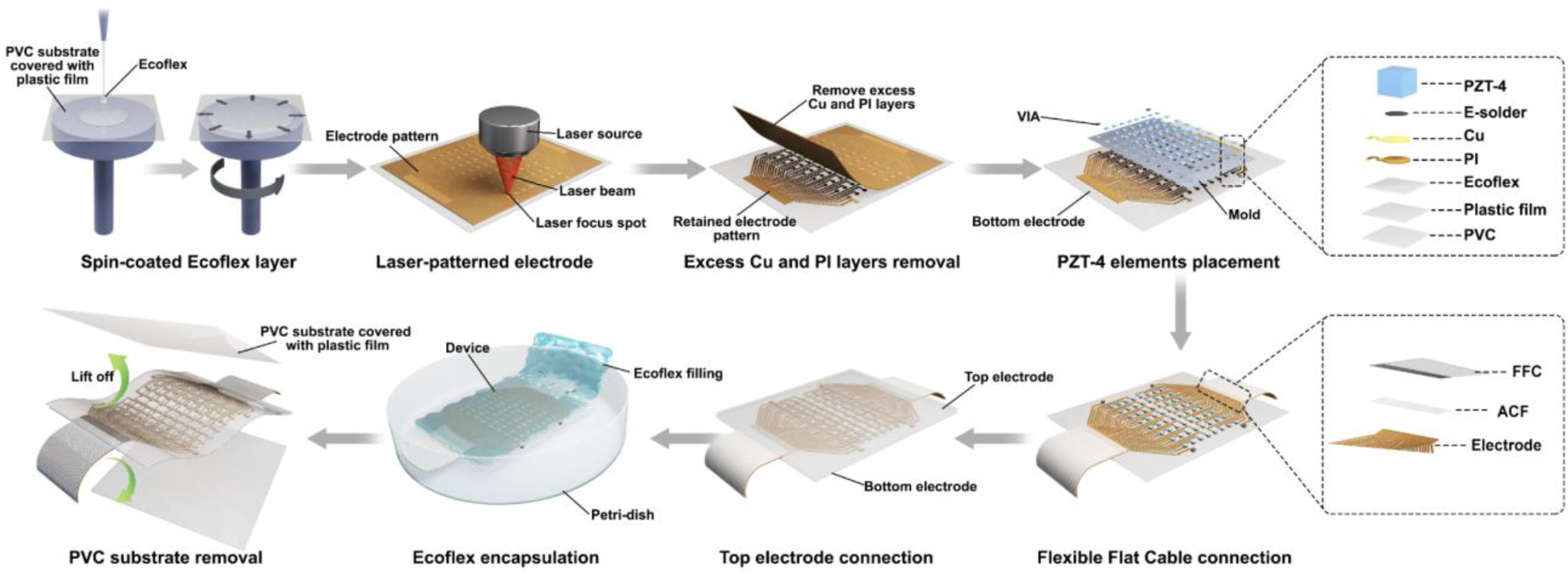
Schematic of the optimized fabrication process for the F-SCAPA.

### 2.2 Mechanical and Electrical Performances

The fabricated F-SCAPA exhibited excellent mechanical performance, with high flexibility enabling natural conformity to various curved surfaces, including a cylindrical surface with a 40 mm diameter (Fig. 3a), a spherical surface with a 120 mm diameter (Fig. 3b), and a human skull (Fig. 3c). Uniaxial tensile testing was performed to evaluate the stretchability of the F-SCAPA, with the setup illustrated in Fig. 3d. Cyclic stress-strain measurements over 20 cycles with 10–50% strain showed consistent stress response, with stress returning to the initial level after each cycle (Fig. 3e). The F-SCAPA sustained a maximum strain of 62.68% with minor stress fluctuations (Fig. 3f), while strains beyond this threshold may cause localized damage to the Ecoflex at the edges of the PZT-4 elements.

**Fig. 3.**
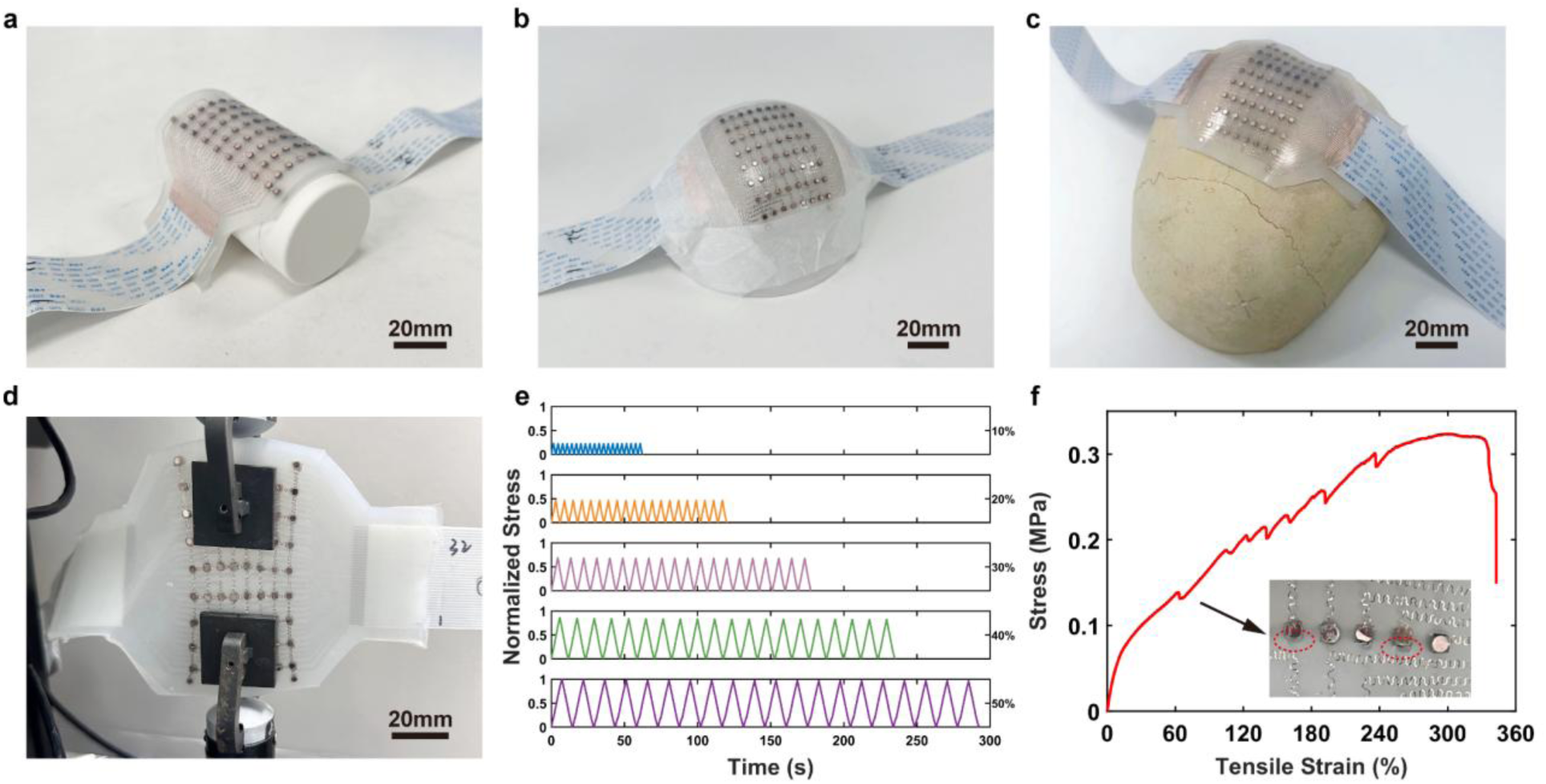
Mechanical characterization of the F-SCAPA. **(a-c)** Photographs of the array conforming to **(a)** a cylindrical surface (40 mm diameter), **(b)** a spherical surface (120 mm diameter), and **(c)** a human skull. **(d)** Optical image of the F-SCAPA under uniaxial stretching. **(e)** Cyclic stress-strain curves demonstrating durability under 20 tensile cycles with 10-50% strain. **(f)** Stress-strain curve under uniaxial tensile loading.

Fig. 4a presents the measured resonance frequencies and corresponding electrical impedance magnitudes for all array elements. Fig. 4b shows the relative impedance change (ΔZ/*Z*_0_ @ *f*_*r*_) under repetitive loading, with the initial impedance *Z*_0_ = 4.01 kΩ. The impedance variations were 0.28%, 0.40%, 0.42%, 0.26%, and 0.16% after 200, 400, 600, 800, and 1000 stretching cycles, respectively. Fig. 4c further illustrates the relative impedance change (ΔZ/*Z*_1_ @ *f*_*r*_) under different stretching ratios, where *Z*_1_ = 4.33 kΩ. At stretching ratios of 10%, 20%, 30%, 40%, and 50%, the impedance changes were 1.92%, −0.29%, −0.76%, 4.11%, and 3.10%, respectively. To overcome the significant impedance mismatch with the 50 Ω driving system, an electrical impedance matching network was implemented to maximize power transfer and minimize reflection loss. After matching, as shown in Fig. 4d, a representative element exhibited a resonant frequency resonant frequency (*f*_*r*_) of 0.502 MHz and an electrical impedance at *f*_*r*_ (Z @ *f*_*r*_) of 38.7 Ω. The key electrical performances across the array are summarized in Fig. 4e-f, with an average resonant frequency (*f*_*r*_) of 0.505 ± 0.003 MHz and average impedance at *f*_*r*_ (Z @ *f*_*r*_) of 40 ± 1.6 Ω.

**Fig. 4.**
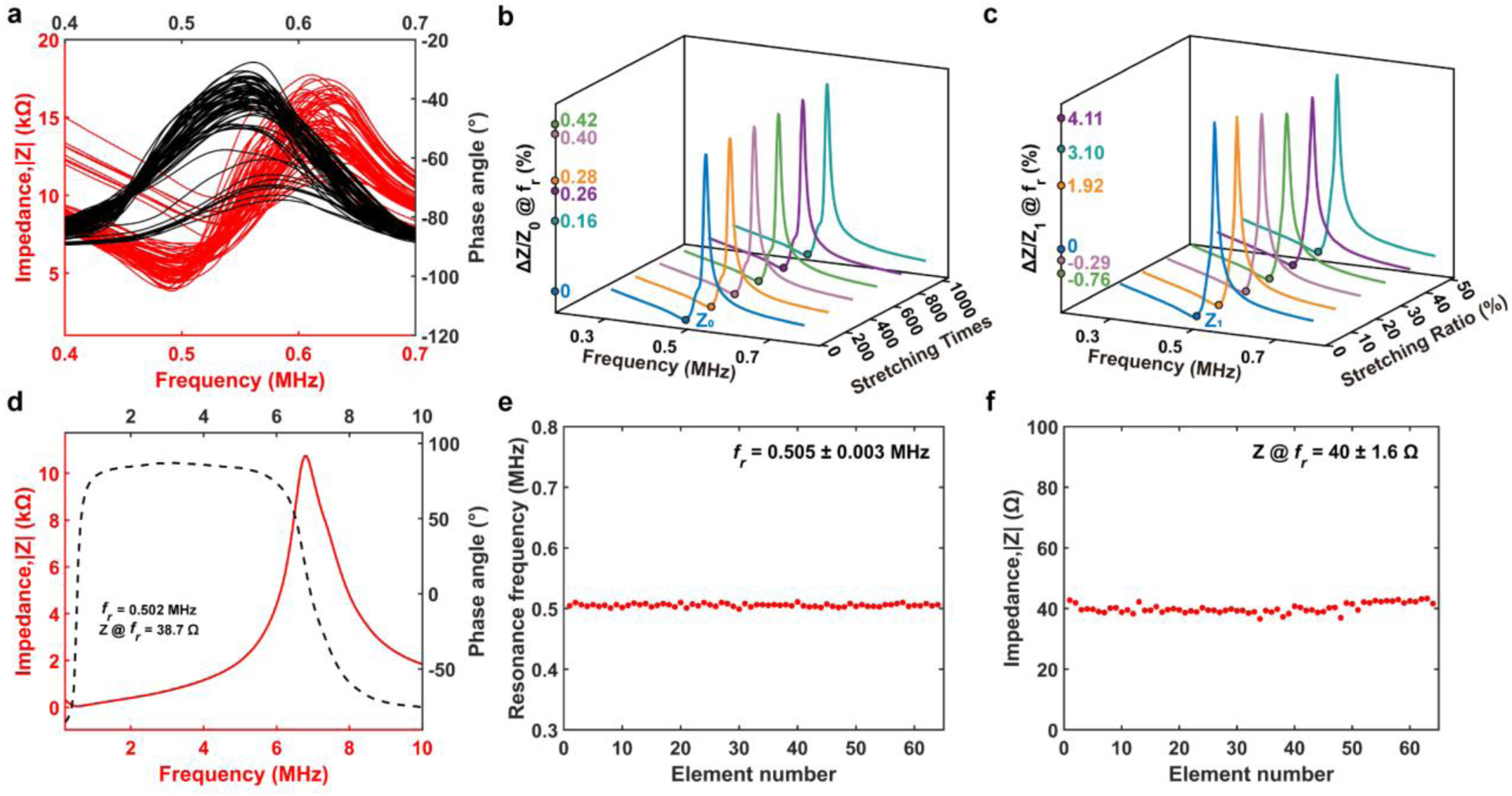
Electrical performance of the F-SCAPA. **(a)** Electrical impedance and phase angle spectra of all array elements. **(b-c)** Relative change in electrical impedance of random array element under **(b)** repetitive loading and **(c)** different stretching ratio. **(d)** Electrical impedance and phase angle spectrum of a representative element after impedance matching. **(e-f)** Uniformity of **(e)** resonance frequency and **(f)** impedance magnitude at response frequency for all elements.

### 2.3 Transcranial Simulation with Aberration Correction

Fig. 5a presents multi-planar CT images and a 3D reconstruction of the skull, providing a detailed model for transcranial simulation with aberration correction that accounts for variations in density, sound speed, and attenuation^29^. The spatial positions of the 64 F-SCAPA elements and three reference markers were captured using a 3D array-camera (Fig. 5b) and then aligned with the CT-derived skull model based on the known marker locations (Fig. 5c).

**Fig. 5.**
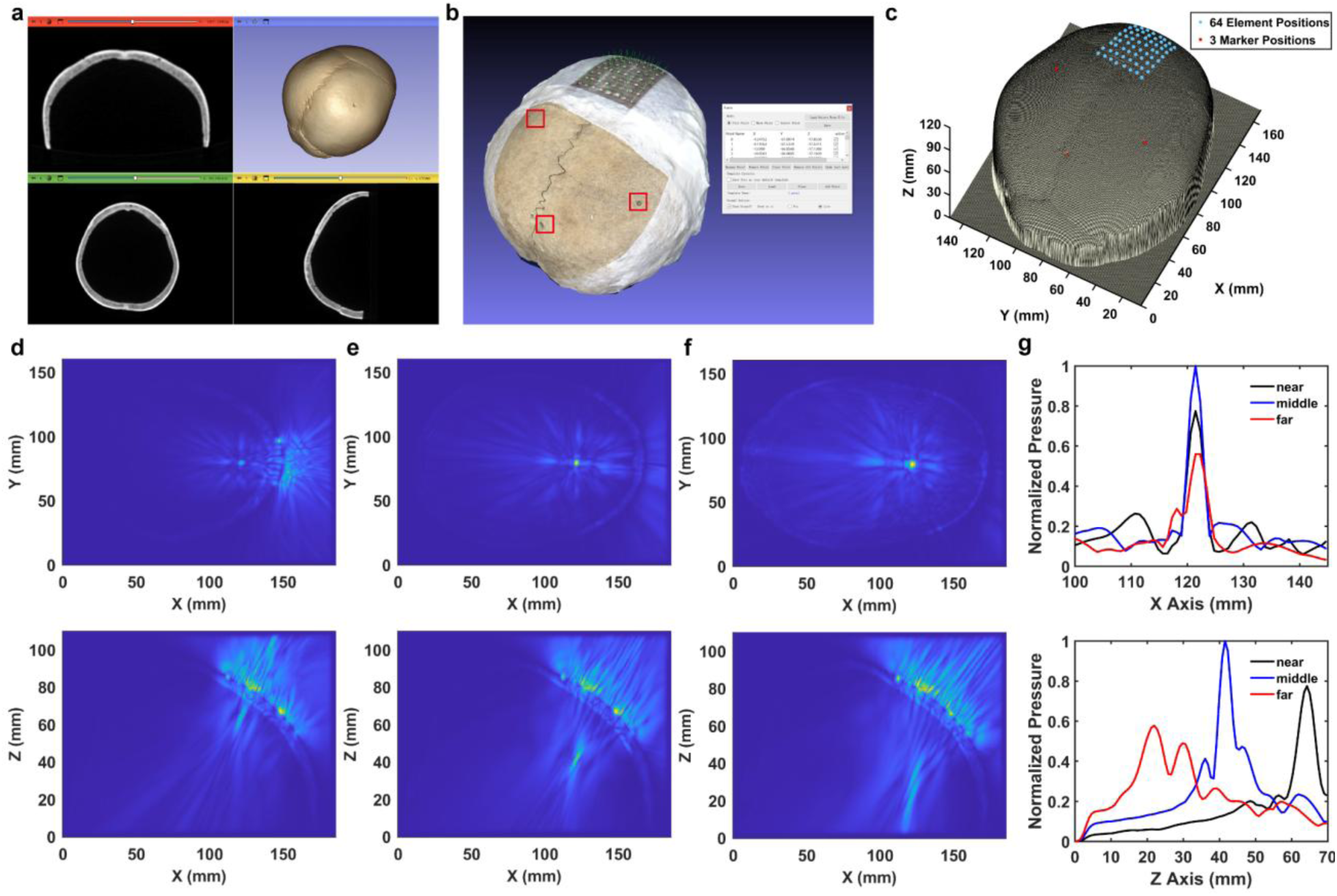
Aberration-Corrected Transcranial Simulation of the F-SCAPA. **(a)** Multi-planar CT image of the human skull. **(b)** Spatial positions of the 64 F-SCAPA elements and three reference markers captured via 3D array-camera scanning. **(c)** Mapping the scanned element and marker positions onto the CT-derived skull model. **(d-f)** Simulated acoustic pressure profiles with aberration correction, focusing at depths of **(d)** 18, **(e)** 36, and **(f)** 50 mm beneath the skull surface. **(g)** Comparison of simulated pressure amplitudes along the X and Z axes at the focus positions.

The time-reversal method was employed to perform simulations with aberration correction, enabling accurate prediction of the acoustic field through the skull. To evaluate the transcranial focusing performance at different depths, three target points were selected at 18 mm (near), 36 mm (middle), and 50 mm (far) beneath the skull surface (Fig. 5d–f). At the near and middle targets, the simulations showed precise focusing, with the maximum pressure located at the intended focal points. At the far target, two adjacent foci appeared near the intended position. The full width at half maximum (FWHM) of the focal spots along the x-axis were 4.99 mm, 4.16 mm, and 4.16 mm, while along the z-axis they were 7.05 mm, 5.48 mm, and 18.02 mm for the near, middle, and far targets, respectively (Fig. 5g).

### 2.4 Experimental Validation of Transcranial Acoustic Fields

After validating the acoustic fields through transcranial simulation, the calculated 64-channel delays were applied on the Vantage HIFU system for experimental transmission. The resulting transcranial acoustic fields are shown in Fig. 6, with FWHM along the x-, y-, and z-directions used to evaluate the beamforming performance of the F-SCAPA. When targeting a focal depth of 36 mm without aberration correction, the acoustic field appeared defocused and scattered (Fig. 6a), with the focal spot shifting to 32.5 mm and exhibiting an elongated z-direction FWHM exceeding 25 mm. After aberration correction, the focal spot was restored to 35.4 mm depth, with FWHM of 5.8, 2.9, and 6.1 mm along the x-, y-, and z-axes, respectively; and an axial FWHM of 8.9 mm (Fig. 6b). At a shallower target of 18 mm, the focal spot exhibited FWHM values of 8.5 mm and 8.2 mm along the x- and y-directions, respectively (Fig. 6c). Fig. 6d summarizes the peak acoustic pressures across these conditions, showing that aberration correction increased the maximum pressure by a factor of 1.30 compared to the uncorrected case.

**Fig. 6.**
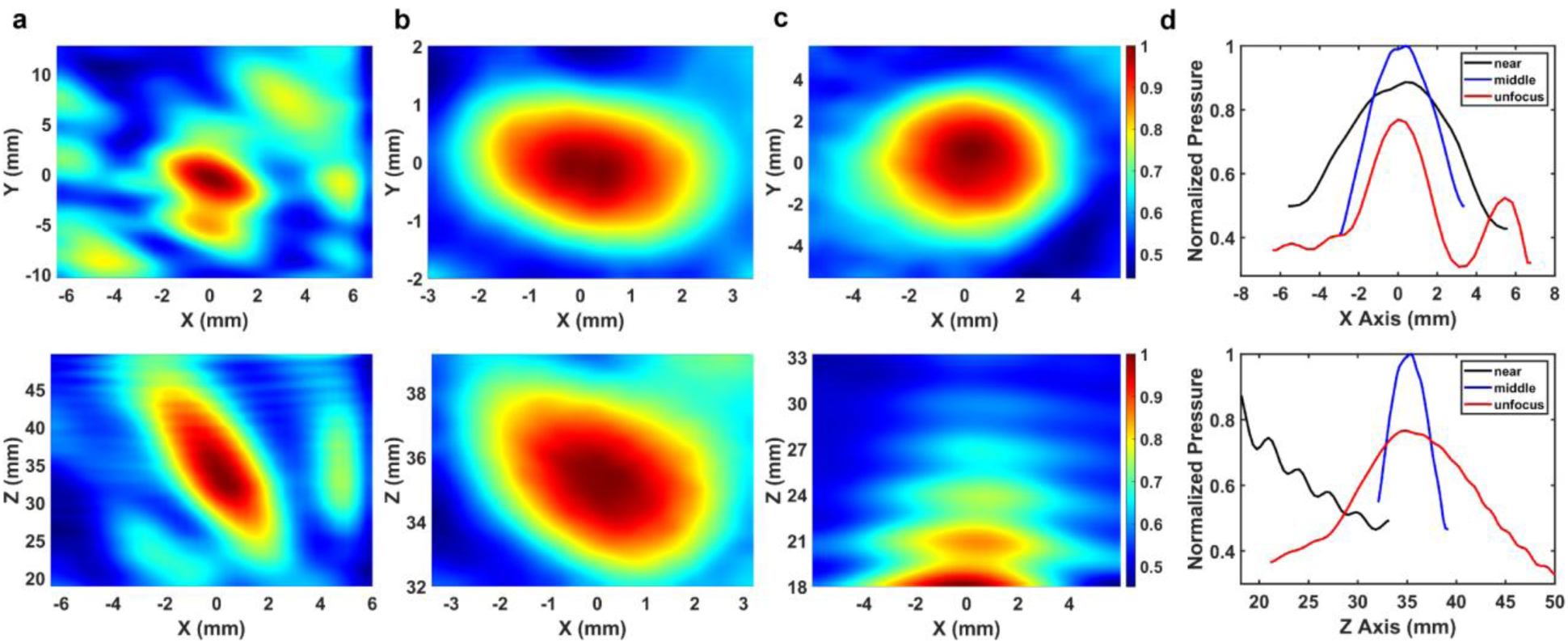
Measured transcranial acoustic pressure distributions at the focus. **(a)** Without aberration correction, the focal spot shifts to 32.5 mm instead of the target 36 mm. **(b)** With aberration correction, the focal spot is restored to 35.4 mm. **(c)** Pressure distribution at a target depth of 18 mm with aberration correction. **(d)** Normalized pressure along the X and Z axes at the focus positions.

## 3. Discussion

This study introduces a flexible skull-conformal aberration-corrected phased array (F-SCAPA) designed for high-precision transcranial ultrasound therapy, overcoming key limitations of rigid systems. By optimizing the array’s structure through Field II simulations, we ensured robust beamforming and deep-brain focusing capabilities. An advanced fabrication process significantly enhanced production efficiency while preserving the array’s mechanical flexibility and acoustic uniformity. Key to our approach is a novel calibration technique that precisely maps element positions on the deformable substrate, enabling accurate integration with transcranial acoustic simulations. This synergy facilitates reliable ultrasound focusing through the skull, addressing challenges of phase aberrations and focal distortion in non-invasive brain interventions.

The performance comparison between the proposed F-SCAPA and previously reported transcranial ultrasound transducers is summarized in Table 1. The F-SCAPA offers several advantages over prior designs. Its conformal geometry improves skull contact and coupling, enabling more stable focusing and reliable phase correction than rigid or bulky structures. Despite operating at 0.5 MHz, it achieves an axial FWHM of 8.9 mm, comparable to rigid arrays at higher frequencies, while supporting a 36 mm focal depth for deeper brain targeting. The fully addressable 8×8 architecture further enables flexible electronic focusing and beamforming. These features highlight the potential of flexible, skull-conformal arrays as a practical platform for non-invasive transcranial ultrasound therapy.

**Table. 1.** Comparison of key parameters of transcranial ultrasound transducers reported in previous studies.

| Research work | Freq. (MHz) | Element number | Structure | Focal distance (mm) | Axial FWHM (mm) | Medium |
| --- | --- | --- | --- | --- | --- | --- |
| Legon et al.<sup>15</sup> | 0.5 | single | Rigid and bulky | 30 | 18 | Human skull |
| Lee et al.<sup>30</sup> | 0.6 | single | Rigid and bulky | 10 | 11.5 | Without skull |
| Manuel et al.<sup>20</sup> | 1 | 128 | Rigid spherical cap | 13 | 9.5 | Human skull |
| Ilham et al.<sup>31</sup> | 0.83 | 16 | Rigid 1D array | 16 | 7.2 | Rat skull |
| Zhang et al.<sup>18</sup> | 1 | 16x16 | Rigid 2D array | 30 | >12 | Without skull |
| **Our work** | **0.5** | **8x8** | **Flexible 2D array** | **36** | **8.9** | **Human skull** |

The F-SCAPA demonstrates superior mechanical flexibility (Fig. 3a-c). Considering that the human skull diameter typically ranges from 8.76 to 19.38 cm^32^, the device was evaluated on a 12 cm curvature model, confirming good conformity to most skull shapes. Uniaxial tensile test revealed a maximum stretchability of 60%, substantially exceeding the maximum tensile strain of human skin (∼20%)^33^, indicating strong adaptability and mechanical durability.

The F-SCAPA exhibits excellent element-to-element uniformity (Fig. 4a), which is essential for precise acoustic focusing. The measured resonance frequency of each element was close to the designed value of 0.5 MHz, with only minor deviations attributed to differences between the actual and simulated material properties. After 1000 stretching cycles, impedance variation remained below 0.42%, and when the stretching ratio increased from 0% to 50%, impedance changes stayed within 4.11%, demonstrating stable electrical performance under mechanical deformation. These slight variations likely result from deformation of the serpentine electrodes and ACF connections, as shown in Fig. 3e, some electrodes are stretched while others are slightly compressed during tension. The impedance matching network effectively reduced the element impedance from approximately 5000 Ω to 40 Ω, closely matching the Vantage system and minimizing energy reflection.

Transcranial simulation results confirmed that the F-SCAPA can successfully focus ultrasound through the skull, with the focal region located close to the intended target positions (Fig. 5d-f). At the middle depth (∼36 mm), the simulated acoustic pressure reached its maximum, and the FWHM was minimal, indicating optimal focusing performance. At the near focus (∼18 mm), the large beamforming angle and reduced geometric phase differences among array elements led to a lower peak pressure and a broader focal spot. At the far focus (∼50 mm), although the time-reversal method compensated for phase aberrations, accumulated errors from CT-based skull modeling, mesh resolution, and element localization occasionally caused focal splitting or slight shifts.

Aberration-corrected simulations showed markedly improved focusing performance, with higher peak pressure and reduced FWHM compared to the uncorrected case (Fig. 6a–b). At a focal depth of 18 mm, hydrophone measurements could not fully capture the entire focal region, but a lower peak pressure and broader focus were still observed. The jittering artifacts in the measured acoustic field were likely caused by reflections at the skull interface. Overall, these results suggest that a focal depth near 35 mm provides optimal transcranial focusing, consistent with many clinical therapeutic applications.

While the results of this study are promising, several areas remain for further improvement. One limitation is the strong acoustic reflections at the skull–transducer interface, mainly due to the large impedance mismatch. This could be addressed by using flexible encapsulation materials with impedance closer to that of the skull to improve acoustic coupling, or by incorporating a targeted waveguide to reduce reflections and guide more energy into the skull. Such strategies would also enhance the F-SCAPA’s ability to focus near the skull surface. In addition, focusing performance at deeper targets could be improved by employing more accurate skull CT models and integrating real-time fMRI feedback to validate and adjust the transcranial simulation model, enabling more precise control and monitoring of the ultrasound focus.

In summary, this study presents an aberration-corrected, flexible 2D phased array for transcranial focused ultrasound therapy, marking the first successful demonstration of a flexible device capable of precise transcranial focusing. The F-SCAPA exhibits excellent mechanical compliance, stable electrical performance, and accurate aberration-corrected acoustic focusing, making it a promising platform for next-generation noninvasive transcranial ultrasound applications.

## 4. Methods

### 4.1 Fabrication Process of the F-SCAPA

The fabrication process began with multiple polyvinyl chloride (PVC) substrates, each covered with thin protective plastic films on both sides. A layer of Ecoflex was spin-coated onto the substrate at 1,000 rpm for 60 s and cured at room temperature for 40 min to form a compliant base layer. A copper-clad polyimide (PI) film (10 µm Cu / 12 µm PI) was then placed on the cured Ecoflex surface, with the copper layer on top. The film was thin enough to conform closely to the Ecoflex. A laser ablation system (LPKF ProtoLaser U4, 1.5 W power, 80 kHz repetition frequency) was used to precisely etch the electrode pattern onto the Cu/PI layer. Optimized laser parameters ensured complete removal of the Cu/PI material in the patterned areas while preserving the Ecoflex layer beneath. Excess Cu/PI in both the top and bottom electrode regions was carefully removed using precision tweezers to finalize the electrode layout.

A mold containing hollow square cavities matching the array layout was fabricated using a high-precision 3D printer (ProJet MJP 2500, 3D Systems, USA) and carefully aligned with the patterned bottom electrode. A bulk PZT-4 sample (3 mm thick) with 5 µm silver electrodes on both surfaces was prepared and diced into 2 mm × 2 mm individual elements. E-Solder 3022 (Von Roll USA, Schenectady, NY) was cured and diced to the same dimensions as the PZT-4 elements to serve as VIA blocks, providing reliable electrical connection between the bottom and top electrodes. A small amount of Conductive adhesive (E-Solder 3022) was dispensed onto each designated element site on the bottom electrode using a fine needle. The mold was then aligned with the electrode, and the PZT-4 elements along with the VIA blocks were precisely positioned into the mold cavities to ensure bonding and electrical connectivity. To avoid adhesion defects, the mold was removed before the E-Solder fully cured (Supplementary Fig. 2).

Electrical connections between the flexible flat cable (FFC) and the bottom electrode were established using anisotropic conductive film (ACF) (Supplementary Fig. 3). The top electrode layer was subsequently bonded to the PZT-4 elements and VIA blocks with Conductive adhesive. After full curing, the entire assembly was encapsulated in Ecoflex and left to cure at room temperature for 2 h. Finally, the PVC substrate and protective film were removed, resulting the fully fabricated F-SCAPA. The detailed acoustic layer stack structure of the F-SCAPA is summarized in Supplementary Table 2.

### 4.2 Basic Performance Characterization

The mechanical properties of the F-SCAPA were characterized using a tensile testing machine (Instron 5944, Instron, USA). The sample was stretched at a constant loading rate of 50 mm/min to evaluate its stretchability and mechanical stability. Cyclic stretching tests were also conducted to examine the repeatability and durability of its mechanical response. The electrical impedance and resonance frequency of each array element were measured using an impedance analyzer (E4990A, Keysight Technologies, USA), with impedance matching achieved through electric impedance matching (EIM) networks (Supplementary Note. 1 and Supplementary Fig.4)^34^. Impedance spectra were measured before and after repeated stretching (200, 400, 600, 800, and 1000 cycles) and under different strain levels (10%, 20%, 30%, 40%, and 50%) to evaluate the electrical stability of the device during mechanical deformation.

### 4.3 Transcranial Aberration-Correction Simulation

A 3D human skull model was reconstructed from CT data with voxel dimensions of 0.47 × 0.47 × 0.50 mm³. The acoustic properties of the skull were derived from the CT Hounsfield units (H) using an established porosity-based empirical model. The porosity map (φ) was computed from the normalized Hounsfield values^35^:

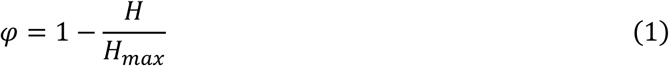

The corresponding mass density (*ρ*), sound speed (c), and attenuation (abs) were then determined as follows^29^:

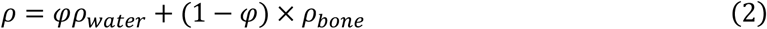

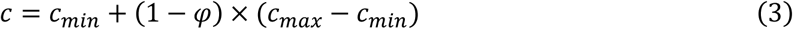

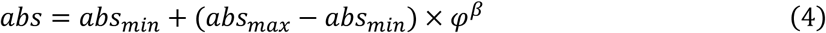

where the optimal agreement between simulation and experimental results was obtained with the following parameters: *ρ*_*water*_ = 1000 *kg*/*m*^3^, *ρ*_bone_ = 2340 *kg*/*m*^3^, *c*_*min*_ = 1540 *m*/*s*, *c*_*max*_ = 2900 *m*/*s*, *abs*_*min*_ = 0.26 *dB*/*mm*, *abs*_*max*_ = 0.87 *dB*/*mm*, *β* = 0.5^36–38^. This mapping generated spatially varying acoustic parameters that reproduced the heterogeneous propagation characteristics of the skull, consistent with prior transcranial ultrasound studies.

Three fiducial markers were identified in both the CT skull model and the 3D array-camera measurements (Super Dimension, Jiaxing, China) to align the two coordinate systems. Based on these markers, the positions of all 64 array elements were first measured in the camera coordinate frame and then mapped to the CT coordinate system using a rigid-body transformation calculated by singular value decomposition (SVD)^39^.

Aberration correction was performed using a time-reversal method implemented in K-Wave. The focal point was modeled as a virtual acoustic source emitting a short pulse (Supplementary Fig. 5) ^40,41^. The back-propagated waves received by all 64 array elements were recorded, and the relative time-of-flight differences were used to calculate individual phase delays. These delays were then applied in the forward simulation to refocus ultrasound through the heterogeneous skull and compensate for phase distortions. Simulations were performed with focal depths of 18, 36, and 50 mm. The resulting acoustic pressure fields were evaluated in the XOY, XOZ, and YOZ planes. The FWHM in the x, y, and z directions was extracted to quantify focusing accuracy and spatial resolution.

### 4.4 Transcranial Acoustic Field Measurement

The F-SCAPA was conformally attached to the curved surface of a human skull (Shanghai Jiao Tong University, School of Medicine, Shanghai, China) and stabilized in degassed water using a custom-designed fixture for acoustic field characterization (Supplementary Fig.6). Phase delay profiles derived from the transcranial simulations were applied to the array via 64 independent channels of the Vantage HIFU system (Vantage 256, Verasonics, USA). To assess the device’s performance, acoustic fields were measured at focal depths of 36 mm (with and without aberration correction) and 18 mm (with aberration correction). A needle hydrophone (NH0200, Precision Acoustics, UK) was used to map the pressure field with a spatial step size of 0.2 mm (Supplementary Fig. 7), and the acoustic signals were recorded using a digital oscilloscope (DSOX3054G, Keysight Technologies, USA).

## Acknowledgements

This work was supported in part by the National Natural Science Foundation of China (Grant No. 12204307).

## Contributions

C. Peng conceived the research idea and directed all research activities. J. Zhang designed the device and performed simulations. J. Zhang fabricated and characterized the device, conducted experiments and wrote the manuscript. H. Qin and C. Pu helped with the device characterization and the experiments. T. Zhang and J. Sun assisted with review, editing and revision of the manuscript. All authors contributed to manuscript writing.

## References

1. Bystritsky, A. & Korb, A. S. A review of low-intensity transcranial focused ultrasound for clinical applications. Curr. Behav. Neurosci. Rep. 2, 60–66 (2015).

2. Azadeh, S. S., Lordifard, P., Soheilifar, M. H., Esmaeeli Djavid, G. & Keshmiri Neghab, H. Ultrasound and sonogenetics: A new perspective for controlling cells with sound. Iran. J. Pharm. Res. IJPR 20, 151–160 (2021).

3. Azadeh, S. S., Lordifard, P., Soheilifar, M. H., Esmaeeli Djavid, G. & Keshmiri Neghab, H. Ultrasound and sonogenetics: A new perspective for controlling cells with sound. Iran. J. Pharm. Res. IJPR 20, 151–160 (2021).

4. Zhu, J., et al. The mechanosensitive ion channel Piezo1 contributes to ultrasound neuromodulation. Proc. Natl. Acad. Sci. 120, e2300291120 (2023).

5. Deng, C. X. Targeted drug delivery across the blood–brain barrier using ultrasound technique. Ther. Deliv. 1, 819–848 (2010).

6. McMahon, D., O’Reilly, M. A. & Hynynen, K. Therapeutic agent delivery across the blood–brain barrier using focused ultrasound. Annu. Rev. Biomed. Eng. 23, 89–113 (2021).

7. Vanneste, S., Reynolds, J. & De Ridder, D. Focused transcranial ultrasound stimulation: A breakthrough approach to treating brain disorders. Expert Rev. Med. Devices 22, 1231–1242 (2025).

8. Luan, S., Williams, I., Nikolic, K. & Constandinou, T. G. Neuromodulation: present and emerging methods. Front. Neuroengineering 7, (2014).

9. Meng, Y., Hynynen, K. & Lipsman, N. Applications of focused ultrasound in the brain: From thermoablation to drug delivery. Nat. Rev. Neurol. 17, 7–22 (2021).

10. Montanaro, H. et al. The impact of CT image parameters and skull heterogeneity modeling on the accuracy of transcranial focused ultrasound simulations. J. Neural Eng. 18, 046041 (2021).

11. Kyriakou, A. et al. A review of numerical and experimental compensation techniques for skull-induced phase aberrations in transcranial focused ultrasound. Int. J. Hyperthermia 30, 36–46 (2014).

12. Xue, X. et al. Flexible ultrasonic transducers for wearable biomedical applications: A review on advanced materials, structural designs, and future prospects. IEEE Trans. Ultrason. Ferroelectr. Freq. Control 71, 786–810 (2024).

13. Bawiec, C. R. et al. A wearable, steerable, transcranial low-intensity focused ultrasound system. J. Ultrasound Med. 44, 239–261 (2025).

14. Yoo, S.-S. Technical Review and Perspectives of Transcranial Focused Ultrasound Brain Stimulation for Neurorehabilitation. Brain Neurorehabilitation 11, (2018).

15. Legon, W. et al. Transcranial focused ultrasound modulates the activity of primary somatosensory cortex in humans. Nat. Neurosci. 17, 322–329 (2014).

16. Kim, H. et al. Miniature ultrasound ring array transducers for transcranial ultrasound neuromodulation of freely-moving small animals. Brain Stimulat. 12, 251–255 (2019).

17. Li, G. et al. Imaging-Guided Dual-Target Neuromodulation of the Mouse Brain Using Array Ultrasound. IEEE Trans. Ultrason. Ferroelectr. Freq. Control 65, 1583–1589 (2018).

18. Zhang, Z. et al. New Sm-PMN-PT Ceramic-Based 2-D Array for Low-Intensity Ultrasound Therapy Application. IEEE Trans. Ultrason. Ferroelectr. Freq. Control 67, 2085–2094 (2020).

19. Adams, C. et al. Implementation of a Skull-Conformal Phased Array for Transcranial Focused Ultrasound Therapy. IEEE Trans. Biomed. Eng. 68, 3457–3468 (2021).

20. Manuel, T. J., Phipps, M. A. & Caskey, C. F. Design of a 1-MHz Therapeutic Ultrasound Array for Small Volume Blood-Brain Barrier Opening at Cortical Targets in Macaques. IEEE Trans. Ultrason. Ferroelectr. Freq. Control 70, 449–459 (2023).

21. Martin, E. et al. Ultrasound system for precise neuromodulation of human deep brain circuits. Nat. Commun. 16, 8024 (2025).

22. Abdullah, A., Shahini, M. & Pak, A. An approach to design a high power piezoelectric ultrasonic transducer. J. Electroceramics 22, 369–382 (2009).

23. Castillo, M., Acevedo, P. & Moreno, E. KLM model for lossy piezoelectric transducers. Ultrasonics 41, 671–679 (2003).

24. Wang, J. et al. Transcranial ultrasound stimulation parameters for neurological diseases: A systematic review. Front. Neurol. 16, (2025).

25. Kubota, J. et al. Real-time monitoring transcranial sub-megahertz ultrasound thrombolysis with phased array scanner. in IEEE Ultrasonics Symposium, 2005. vol. 3 1716–1719 (2005).

26. Zhang, T., Pan, N., Wang, Y., Liu, C. & Hu, S. Transcranial Focused Ultrasound Neuromodulation: A Review of the Excitatory and Inhibitory Effects on Brain Activity in Human and Animals. Front. Hum. Neurosci. 15, 749162 (2021).

27. Li, P.-C. & Huang, J.-J. Efficient dynamic focus control for three-dimensional imaging using two-dimensional arrays. IEEE Trans. Ultrason. Ferroelectr. Freq. Control 49, 1191–1202 (2002).

28. di Biase, L., Falato, E. & Di Lazzaro, V. Transcranial Focused Ultrasound (tFUS) and Transcranial Unfocused Ultrasound (tUS) Neuromodulation: From Theoretical Principles to Stimulation Practices. Front. Neurol. 10, (2019).

29. Aubry, J. F., Tanter, M., Pernot, M., Thomas, J. L. & Fink, M. Experimental demonstration of noninvasive transskull adaptive focusing based on prior computed tomography scans. J. Acoust. Soc. Am. 113, 84–93 (2003).

30. Lee, W. et al. Transcranial focused ultrasound stimulation of motor cortical areas in freely-moving awake rats. BMC Neurosci. 19, 57 (2018).

31. Ilham, S. J., Kashani, Z. & Kiani, M. Design and Optimization of Ultrasound Phased Arrays for Large-Scale Ultrasound Neuromodulation. IEEE Trans. Biomed. Circuits Syst. 15, 1454–1466 (2021).

32. Pirouzmand, F. & Muhajarine, N. Definition of topographic organization of skull profile in normal population and its implications on the role of sutures in skull morphology. J. Craniofac. Surg. 19, 27 (2008).

33. Liu, H. et al. Flexible ultrasonic transducer array with bulk PZT for adjuvant treatment of bone injury. Sensors 20, 86 (2020).

34. Moon, J., Lee, J. & Chang, J. H. Electrical impedance matching networks based on filter structures for high frequency ultrasound transducers. Sens. Actuators Phys. 251, 225–233 (2016).

35. Guo, S. et al. Feasibility of ultrashort echo time images using full-wave acoustic and thermal modeling for transcranial MRI-guided focused ultrasound (tcMRgFUS) planning. Phys. Med. Biol. 64, 095008 (2019).

36. Aubry, J. F., Tanter, M., Pernot, M., Thomas, J. L. & Fink, M. Experimental demonstration of noninvasive transskull adaptive focusing based on prior computed tomography scans. J. Acoust. Soc. Am. 113, 84–93 (2003).

37. Webb, T. D. et al. Acoustic attenuation: Multifrequency measurement and relationship to CT and MR imaging. IEEE Trans. Ultrason. Ferroelectr. Freq. Control 68, 1532–1545 (2021).

38. Hüter, Th. F. Messung der ultraschallabsorption im menschlichen schädelknochen und ihre abhängigkeit von der frequenz. Naturwissenschaften 39, 21–22 (1952).

39. Challis, J. H. A procedure for determining rigid body transformation parameters. J. Biomech. 28, 733–737 (1995).

40. Thomas, J.-L. & Fink, M. A. Ultrasonic beam focusing through tissue inhomogeneities with a time reversal mirror: Application to transskull therapy. IEEE Trans. Ultrason. Ferroelectr. Freq. Control 43, 1122–1129 (1996).

41. Treeby, B. E. & Cox, B. T. k-wave: MATLAB toolbox for the simulation and reconstruction of photoacoustic wave fields. J. Biomed. Opt. 15, 021314 (2010).

