## Supplementary information for "Flexible Skull-Conformal Phased Array for Aberration-Corrected Transcranial Focused Ultrasound Therapy"

### Table of Contents

**Supplementary Note. 1 Principle of Electric Impedance Matching**

**Supplementary Tables. 1-2**

**Supplementary Figures. 1-7**

**Supplementary References**

#### Supplementary Note. 1 Principle of Electric Impedance Matching

The flexible skull-conformal aberration-corrected phased array (F-SCAPA) exhibits an impedance significantly higher than the standard 50 Ω of the driving system. This impedance mismatch can result in poor energy transfer and reduced transmitting efficiency. To overcome this, a series–parallel L-type electric impedance matching (EIM) network is employed, maximizing power delivery and optimizing transducer performance (**Supplementary Figure. 4**). The network consists of a series inductor $\mathrm{L_s}$ and a parallel capacitor $\mathrm{C_p}$. Their values are determined based on the transducer impedance, $Z_x = R_x + jX_x$, and the system impedance, $Z_s = R_s + jX_s$, at the operating frequency $f$, according to the following equations[1]:

$$\mathrm{L_s} = \frac{-R_x X_s + \sqrt{R_x R_s [R_x^2 + X_x^2 - R_x R_s]}}{2\pi f \cdot R_x} \tag{1}$$

$$\mathrm{C_p} = \frac{-X_x R_s + \sqrt{R_x R_s [R_x^2 + X_x^2 - R_x R_s]}}{2\pi f \cdot (R_s - R_x)} \tag{2}$$

Here, $R_x$ and $X_x$ are the real and imaginary parts of the transducer impedance, and $R_s$ and $X_s$ are the real and imaginary parts of the system impedance. At resonance, the reactive components of the F-SCAPA and system cancel each other, resulting in an effective impedance closely matching the system. This configuration ensures maximum power transfer in accordance with the maximum power transfer theorem. Implementation of the EIM network reduces reflection losses and improves acoustic output.

**Supplementary Table. 1 Comparison of parameters of different types of piezoelectric materials.**

| Properties | Unit | PZT-4[2,3] | PZT-5H[4] | 1-3 PZT 5H Composite[5,6] |
|---|---|---|---|---|
| Mechanical Quality Factor, Qm | - | 600 | 70 | 88 |
| Piezoelectric charge coefficient, $d_{33}$ | pC/N | 280 | 650 | 593 |
| Electromechanical coupling coefficient, $k_t$ | - | 0.55 | 0.46 | 0.55 |
| Curie Temperature, $T_c$ | °C | 320 | 200 | 180 |
| Density, ρ | $Kg/m^3$ | 7500 | 7750 | 4600 |
| Longitudinal velocity, v | m/s | 4630 | 4560 | 3600 |

**Supplementary Table. 2 Acoustic layer stack structure of the F-SCAPA.**

| Acoustic layer | Materials | Thickness |
|---|---|---|
| Package layer | Ecoflex | 15 μm |
| Electrode | PI | 12 μm |
| | Cu | 10 μm |
| Bonding layer | E-solder | 5 μm |
| Piezoelectric layer | PZT-4 | 3 mm |
| Bonding layer | E-solder | 5 μm |
| Electrode | Cu | 10 μm |
| | PI | 12 μm |
| Package layer | Ecoflex | 15 μm |

**Supplementary Figure. 1**

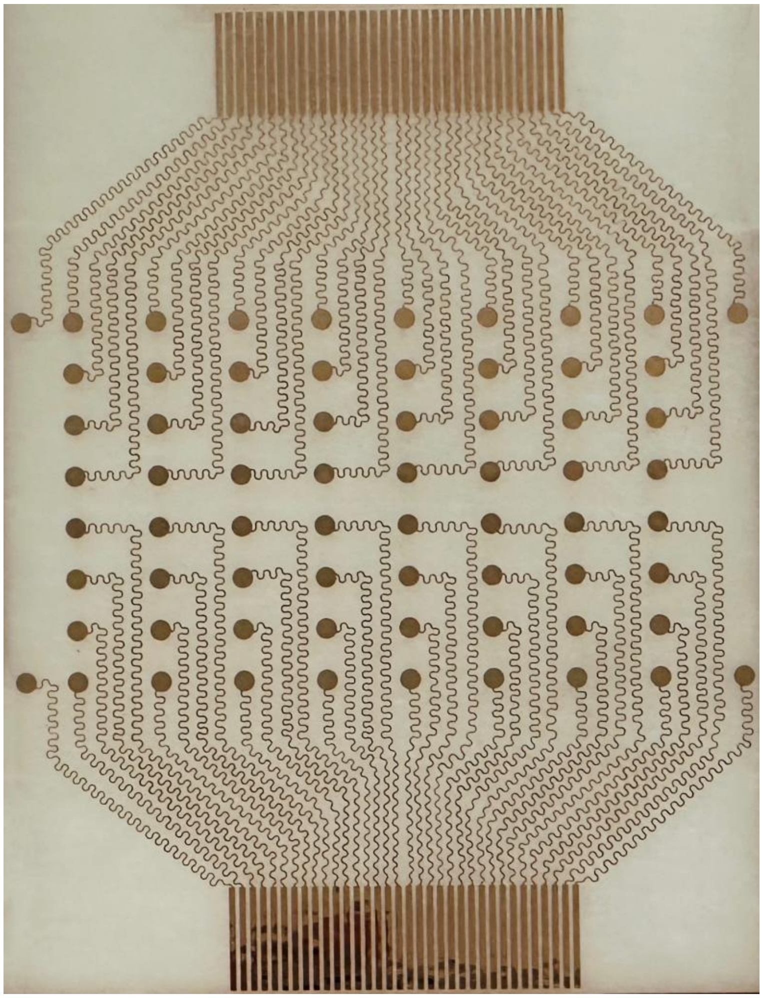

**Supplementary Fig. 1** Electrode pattern on Ecoflex-coated PVC substrate.

**Supplementary Figure. 2**

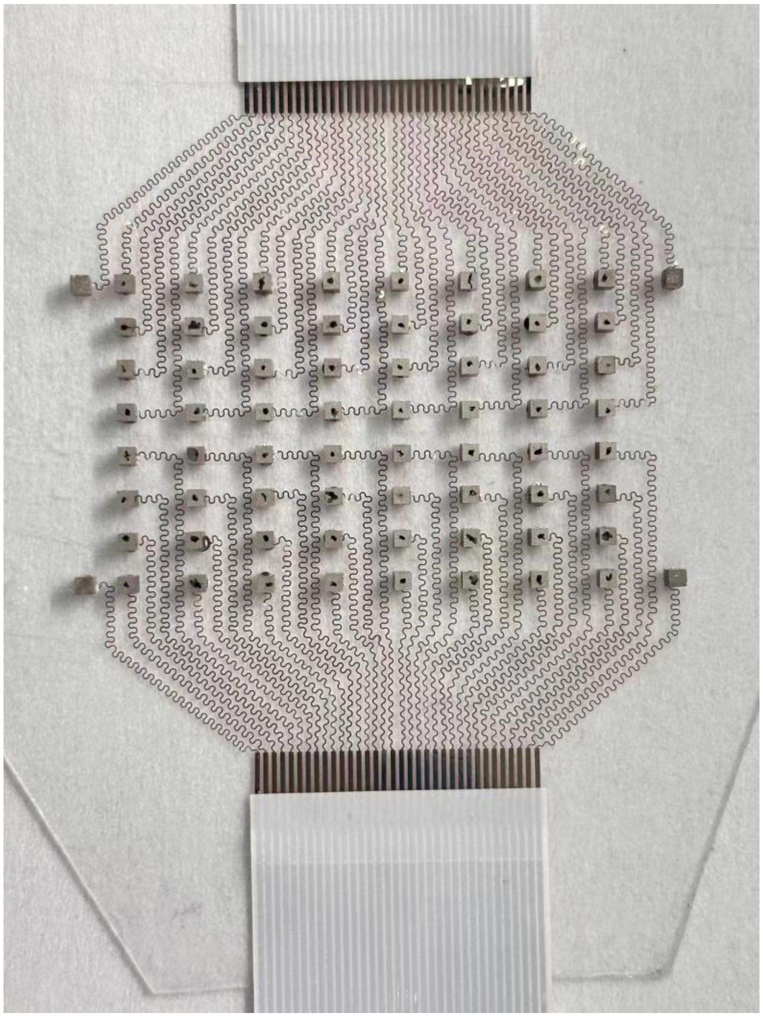

**Supplementary Fig. 2** PZT-4 elements aligned on electrodes following array layout after mold removal.

**Supplementary Figure. 3**

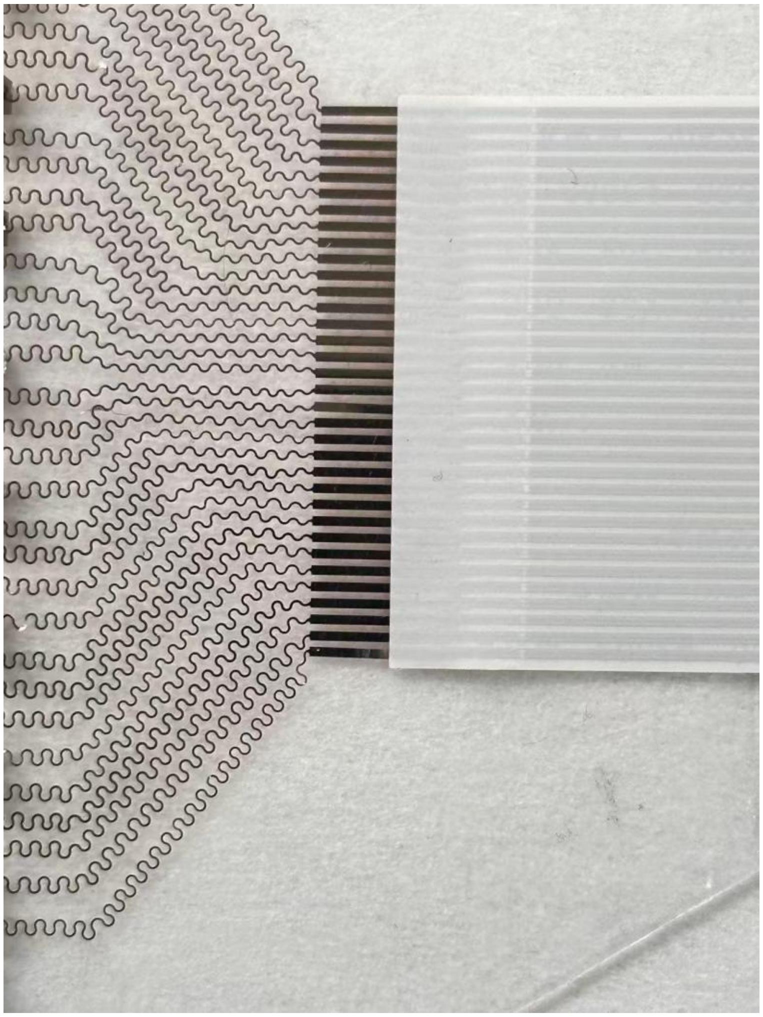

**Supplementary Fig. 3** Electrical connection between the flexible flat cable (FFC) and bottom electrode using anisotropic conductive film (ACF).

**Supplementary Figure. 4**

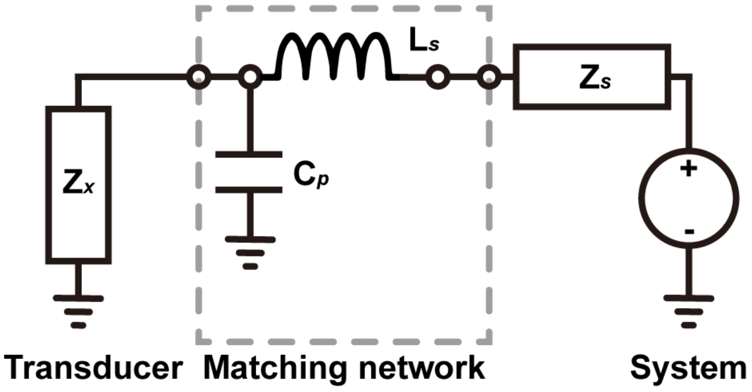


**Supplementary Fig. 4** Schematic block diagram of the electric impedance matching (EIM) networks.

**Supplementary Figure. 5**

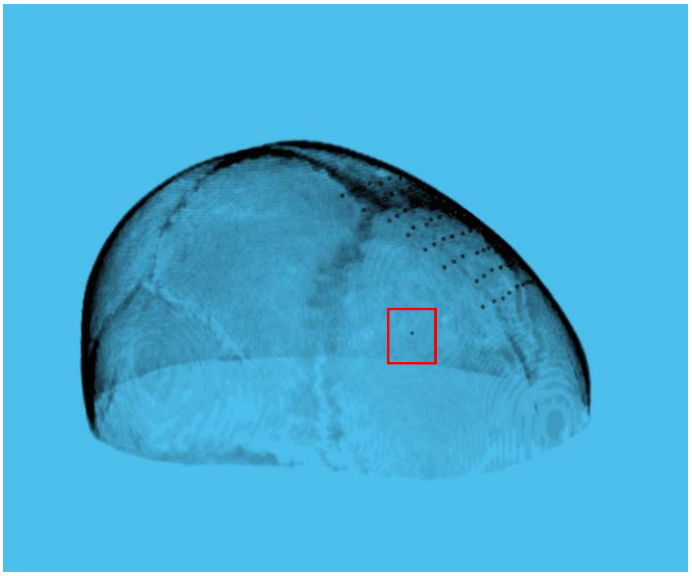

**Supplementary Fig. 5** Illustration of the virtual source in K-Wave time-reversal simulation.

**Supplementary Figure. 6**

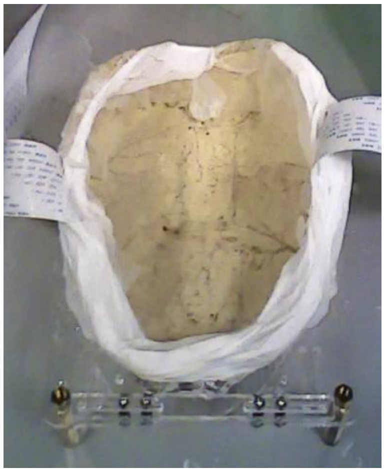

**Supplementary Fig. 6** Experimental fixture with F-SCAPA for transcranial measurements.

**Supplementary Figure. 7**

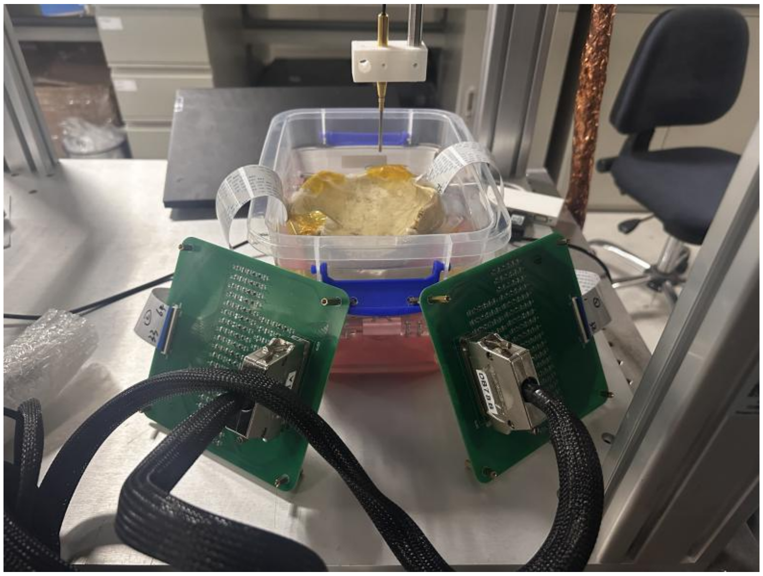

**Supplementary Fig. 7** Experimental setup for transcranial acoustic field measurement with the F-SCAPA.

#### Supplementary References


1. Moon, J., Lee, J. & Chang, J. H. Electrical impedance matching networks based on filter structures for high frequency ultrasound transducers. Sens. Actuators Phys. 251, 225–233 (2016).
2. Abdullah, A., Shahini, M. & Pak, A. An approach to design a high power piezoelectric ultrasonic transducer. J. Electroceramics 22, 369–382 (2009).
3. Castillo, M., Acevedo, P. & Moreno, E. KLM model for lossy piezoelectric transducers. Ultrasonics 41, 671–679 (2003).
4. Peng, C., Chen, M. & Jiang, X. Under-display ultrasonic fingerprint recognition with finger vessel imaging. IEEE Sens. J. 21, 7412–7419 (2021).
5. Li, L. et al. 1–3 piezoelectric composites for high-temperature transducer applications. J. Phys. Appl. Phys. 46, 165306 (2013).
6. Kim, K., Hsu, D. K., Ahn, B., Kim, Y.-G. & Barnard, D. J. Fabrication and comparison of PMN-PT single crystal, PZT and PZT-based 1-3 composite ultrasonic transducers for NDE applications. Ultrasonics 50, 790–797 (2010).
